# The effect of age and sex hormones on female neuromuscular function across the adult lifespan

**DOI:** 10.1101/2024.11.29.626123

**Authors:** Steven J. O’Bryan, Annabel Critchlow, Cas Fuchs, Danielle Hiam, Séverine Lamon

## Abstract

Neuromuscular ageing is characterized by neural and/or skeletal muscle degeneration that decreases maximal force and power. Female neuromuscular ageing occurs earlier in life compared to males, potentially due to sex hormone changes during the menopausal transition. We quantified neuromuscular function in 88 healthy females represented equally over each decade from 18-80 years of age and investigated the potential role of decreased ovarian hormone concentrations following menopause. Neuromuscular assessment included quadriceps maximal voluntary and evoked isometric torque and surface electromyography measurements, plus one-repetition maximum leg press. Voluntary and evoked torques and one-repetition maximum decreased non-linearly with age, with accelerated reductions starting during the fourth decade. An absence of changes in volitional recruitment of existing quadriceps motor units and Ia afferent facilitation of spinal motoneurons suggests that functional decline was largely mediated by impairment in intrinsic muscle function and/or neuromuscular transmission. Maximal muscle compound action potential amplitude decreased with increasing age for rectus femoris only, indicating increased vulnerability to neuromuscular degeneration compared to vastus lateralis and medialis. In postmenopausal females, some variance in data was explained by inter-individual differences in body composition and physical activity level, however, changes in total or free concentrations of oestrogen, progesterone and/or testosterone were correlated with all age-related decreases in neuromuscular variables. In conclusion, we demonstrate an accelerated onset of neuromuscular degeneration of muscular origin around menopause onset, which is associated with changes in sex hormone concentrations. Interventions aimed at mitigating declines in ovarian hormones and their subsequent effects on neuromuscular function postmenopause should be further explored.

**Key Points:**

- Neuromuscular deterioration with age is associated with poor physical function and quality of life in older adults, but female-specific trajectories and mechanisms remain unclear.
- This study is the first to map neuromuscular function across each decade of the adult lifespan in 88 healthy females from 18 to 80 years old and to examine the potential role of hormonal changes after menopause
- We show an accelerated reduction in neuromuscular function, primarily of muscular origin, that occurs between 40 and 50 years of age, coinciding with the onset of menopause.
- In postmenopausal females, age-related reductions in neuromuscular function can be explained by differences in body composition, physical activity, and sex hormone concentrations.
- These findings help us better understand the factors that contribute to the loss of neuromuscular function with age in females, enabling the identification of potential therapeutic interventions to prolong the female health span.

## Introduction

Neuromuscular function describes the integration and translation of multiple synaptic potentials generated and received by upper and lower motoneurons into force or torques produced by the musculotendinous unit, thereby encapsulating mechanisms spanning the entire motor pathway. Ageing has detrimental effects on neuromuscular function characterized by a decrease in maximal muscle strength or power, which may be associated with reduced motor recruitment or loss in skeletal muscle mass and quality (Hunter *et al*., 2016; Tieland *et al*., 2018; O’Bryan & Hiam, 2022). For example, isometric muscle strength declines by ∼1 - 5% per year between ∼ 60 – 80 years of age (Hughes *et al*., 2001; Delmonico *et al*., 2009; Kim *et al*., 2018) with older males and females greater than 60 years of age being ∼20-40% weaker than their younger counterparts (Lindle *et al*., 1997; Wu *et al*., 2016; Roberts *et al*., 2018). The decline in strength and power occurs earlier in the lifespan for females when compared to males (Lindle *et al*., 1997; Roberts *et al*., 2018) and is accelerated at a faster rate beyond 80 years of age (Kim *et al*., 2018). Moreover, females exhibit a higher frailty index (Gordon *et al*., 2017) leading to a larger population of older females accessing aged-care facilities and requiring assistance with activities of daily living (Austad, 2006).

Despite the evidence that females display different trajectories in the age-related decline in neuromuscular function compared to males, the mechanisms responsible for this dimorphic response remain unclear largely due to the inherent underrepresentation of females in physiological research (Garcia-Sifuentes & Maney, 2021). Distinct transient fluctuations in the synthesis and concentration of primary ovarian female sex-hormones oestrogen and progesterone, but also testosterone, across the different phases of the female lifespan including premenopausal (∼18 – 45 years), perimenopausal (∼40-55 years) and postmenopausal (∼50 years onward) periods is one primary potential cause (Hansen & Kjaer, 2014; Alexander *et al*., 2022; Critchlow *et al*., 2023; Piasecki *et al*., 2024). Indeed, sex hormone-sensitive nucleic or membrane-bound receptors are found throughout both the central nervous (Barth *et al*., 2015) and musculoskeletal systems (Wiik *et al*., 2009). Animal models demonstrate a clear effect of steroidal sex hormones (or their precursors) on neuromuscular function (Callachan *et al*., 1987; Smith *et al*., 2002; Moran *et al*., 2007; Schultz *et al*., 2009; Barth *et al*., 2015; Collins *et al*., 2019; Piasecki *et al*., 2024). In ovariectomized female rats administration of progesterone potentiates gamma-aminobutyric acid (GABA) inhibition and suppresses glutamate-induced excitation (Smith *et al*., 1987). Conversely, oestrogen suppresses GABAergic neurons and attenuates the release of GABA in pyramidal cells (Schultz *et al*., 2009), enhances glutamatergic transmission (Barth *et al*., 2015), and possesses potential excitatory effects modulated by dopaminergic (Rey *et al*., 2014) and serotonergic pathways (Lu *et al*., 1999). Beyond the neuromuscular junction, ovariectomized mice with oestrogen deficiency demonstrate reduced skeletal muscle contractility (Moran *et al*., 2007), cross-sectional area (Moran *et al*., 2007) and satellite cell number in fast-twitch skeletal muscles (Collins *et al*., 2019) that can be restored with oestradiol treatment. Although cause and effect relationships in humans are more difficult to define, female ageing studies provide an observational model to gain understanding of the role of primary sex hormones on neuromuscular function. Indeed, during perimenopause and postmenopause, oestradiol reduces to one-third and one-fifth of that observed during the reproductive years, respectively, whereas progesterone reduces to two-thirds the level of younger females during perimenopause and is essentially absent in postmenopause (Landgren *et al*., 2004; Burger *et al*., 2008). The potential role female sex hormones play in the observed decline in neuromuscular function in older females has however not been investigated in a large cohort of participants.

Quadriceps strength and power is associated with functional capability throughout the lifespan (Fuchs *et al*., 2023) leading several studies to investigate the effects of ageing and the menopausal transition on quadriceps neuromuscular function in females. Compared to young females, postmenopausal females less than 65 years of age generate lower (Pöllänen *et al*., 2011; Pöllänen *et al*., 2015) or comparable (Laakkonen *et al*., 2017) absolute or specific (% cross-sectional area) isometric torque during quadriceps maximal voluntary contraction (MVC), with similar conflicting results reported for quadriceps muscle cross sectional area (CSA) (Ahtiainen *et al*., 2012; Pöllänen *et al*., 2015; Collins *et al*., 2019; Park *et al*., 2019; Juppi *et al*., 2020; Pesonen *et al*., 2021). However, in postmenopausal females beyond the age of 65 years, lower quadriceps MVC and CSA compared to younger females is more consistently reported (Häkkinen *et al*., 1996; Wu *et al*., 2016; Yacyshyn & McNeil, 2020; Varesco *et al*., 2022; Wrucke *et al*., 2024) and may be accompanied by reduced neural drive and voluntary activation (Mau-Moeller *et al*., 2013; Wu *et al*., 2016; Solianik *et al*., 2017; Rozand *et al*., 2020), rate of torque development (Häkkinen *et al*., 1996; Yacyshyn & McNeil, 2020; Wrucke *et al*., 2024) and potentiated evoked torques (Solianik *et al*., 2017; Yacyshyn & McNeil, 2020; Varesco *et al*., 2022; Wrucke *et al*., 2024). However, many comparison studies between younger and older female cohorts are limited in their capacity to define the trajectory of neuromuscular changes across the lifespan and to investigate the critical perimenopausal period, which is marked by severe unpredictable and fluctuating hormonal patterns (Landgren *et al*., 2004; Burger *et al*., 2007). Of the limited studies describing neuromuscular changes during the perimenopausal phase, subtle within-phase changes in quadriceps isometric MVC and evoked twitch torque have been suggested (Pesonen *et al*., 2021). Larger cohort studies and scoping reviews of a wider age range of females covering each decade of the lifespan (20-80 years of age) suggest an accelerated decline in quadriceps MVC occurring around perimenopause (Lindle *et al*., 1997; Haynes *et al*., 2020) that may (Hughes *et al*., 2001; Mizuno *et al*., 2021) or may not (Rolland *et al*., 2007; Delmonico *et al*., 2009) be associated with reduced skeletal muscle mass. Thus, functional decline occurring during the perimenopause period may also be explained by several neuromuscular factors including alterations in motor unit recruitment (Piasecki *et al*., 2024), sarcolemmal excitability (Lee *et al*., 2018) and/or excitation-contraction coupling including Ca^2+^ mediated cross-bridge formation (Mazara *et al*., 2021). Currently, age-related trajectories of neuromuscular dysfunction across the female lifespan and including premenopausal, perimenopausal and postmenopausal periods remains to be thoroughly investigated.

The aims of this study were to quantify age-related changes in quadriceps neuromuscular function across all phases of the female lifespan and to investigate how inter-individual differences in body composition, physical activity level and primary sex hormones (oestrogen, progesterone and testosterone) contribute to any observed decline postmenopause. To address this aim, voluntary and evoked isometric quadriceps torque and surface electromyography outcomes, one-repetition maximum leg press, body composition including isolated quadriceps lean cross-sectional area, and female sex hormone concentrations were measured in 88 apparently healthy females 18-80 years of age.

## Method

### Ethical approval and Participants

The present study was approved by Deakin University’s Human Research Ethics Committee (DUHREC 2021-307) and conducted in accordance with the standards set by the Declaration of Helsinki. All volunteers provided written informed consent after being provided with a digital copy of the plain language summary of the study, which detailed the experimental procedures, associated risks, and the liberty to withdraw consent at any time without jeopardy. Eligibility for inclusion included apparently healthy biological females between 18-80 years of age. Health status was determined via a medical history questionnaire and exclusion criteria included pregnancy, cancer, implanted medical devices, BMI > 35kg.m^2^ or musculoskeletal injury/pain of the tibiofemoral or patellofemoral joint complex. Eighty-eight participants (age range 19 – 77 years) took part in this study. Participants were instructed to avoid strenuous physical activity in day before and day of neuromuscular assessments and to avoid caffeine on the day.

### Neuromuscular assessment of the quadriceps

Specific information pertaining to quadriceps neuromuscular assessment methodology including electrical stimulation, surface electromyography and data acquisition and analysis have all been described in detail by our group previously (O’Bryan *et al*., 2024). Briefly, isometric neuromuscular assessments were conducted on an isokinetic dynamometer (Universal Pro Single Chair model 850-230, Biodex Medical Systems, United States). Participants sat upright in the dynamometer chair with hip and knee angle set at 85° and 75° flexion, respectively. Participants began each neuromuscular assessment with a standardized warm-up procedure including a series of voluntary submaximal (20%, 40%, 60%, 80% perceived effort) and minimum of three maximal (100% perceived effort) isometric knee extensions and knee flexions (4s push, 1s relax, 4s pull) until a plateau in voluntary maximal torque was recorded (60s between maximal efforts). Following the warm-up, participants performed a ⁓4s maximal isometric voluntary contraction (MVC) of the knee extensors with an electrically evoked doublet (100_Hz_) applied to the femoral nerve at the plateau in voluntary torque, followed ∼2s after by three resting evoked twitch responses (100_Hz_, 10_Hz_ and 1_Hz_ ∼1.5s apart). This procedure was repeated three times (two minutes rest separating each set) with the highest recorded MVC (N·m), quadriceps root mean square normalized to maximal compound muscle action potential (RMS.M_MAX_), voluntary activation (VA %) peak potentiated resting twitch torque (100_Hz_ and 10_Hz_ in N·m plus a 100:10_Hz_ ratio), maximal rate of torque development (N·m.s^-1^) and quadriceps maximal compound muscle action potential peak-to-peak amplitude (M_MAX_) and duration (M_DUR_) calculated for analysis. After ∼15 min rest period, Hoffman reflexes (H-reflex) were elicited in quadriceps by applying a 1_Hz_ electrical stimulus to the femoral nerve at progressively increasing intensity during 50 brief (2-3s) submaximal isometric contractions performed at 5% MVC (10s rest separated contractions), with the highest peak-to-peak amplitude of the H-reflex normalized to the peak-to-peak amplitude of the maximal compound muscle action potential (H_MAX_.M_MAX_) included for analysis. Finally, after another ∼15 min rest period, dynamic leg strength was assessed with a leg-press 5-repetition maximum (RM) test. Participants completed up to four test sets of five repetitions. After each test set, participants were asked to rate their perceived exertion from 1 to 10 on the modified Borg Scale (Borg, 1982). The resistance was increased until the participant communicated a 10 rating (i.e., maximally exerted) after five repetitions. The corresponding weight was recorded as their 5RM (kg), and a modified equation was used to calculate estimated one-repetition maximum leg press (e1RM in kg) (Wood *et al*., 2002):

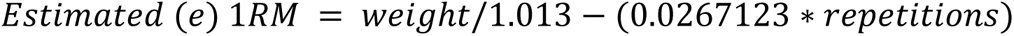

The first twenty-five participants completed two neuromuscular assessment sessions (minus 5RM test) with excellent inter-session reliability established for all isometric voluntary and evoked outcomes (O’Bryan *et al*., 2024). Accordingly, the remaining sixty-three participants were assessed once. Herein, voluntary and evoked isometric torques and e1RM were additionally reported relative to quadriceps lean muscle cross-sectional area (N·m.mm²) to describe muscle-specific torque and quality (Pöllänen *et al*., 2011).

### Physical activity status

Participants recorded their physical activity across a 7-day period with an actiCAL accelerometer watch (Respironics Inc, Murraysville, PA). This watch records energy expenditure and intensity of physical activity and has been validated against oxygen consumption-computed active energy expenditure (Heil, 2006). During the hours in which participants were awake (determined by a sleep diary) the time spent in moderate and vigorous physical activity (>1,535 counts per minute; MVPA) per day was calculated and averaged across all valid days (Colley & Tremblay, 2011). Days were considered valid if there was >10 hours of data available, and participants required at least 4 valid days of data to be included in the physical activity analysis. Eight participants did not meet these criteria and were excluded from physical activity analysis.

### Hormonal Status

Fasted plasma samples were taken from the antecubital vein, immediately centrifuged at 4°C, and stored at -80°C until analysed. Oestradiol, testosterone, progesterone, luteinising hormone (LH), and follicle-stimulating hormone (FSH) were measured by high-performance liquid chromatography mass spectrometry (Triple Quad 5500, Sciex, Framingham, MA) at Monash Health Pathology laboratory (Victoria, Australia). Conjointly, a menstrual cycle history questionnaire was used to determine self-reported menopausal stage as well as use of hormonal contraception or menopausal hormone replacement therapy. Information was also collected about the last known menstrual cycle, reproductive conditions (polycystic ovary syndrome: n=5, endometriosis n=2, fibroids: n=5, or ovarian cysts: n=5) and symptoms of perimenopause or menopause (irregular periods, muscle/joint pain, sleep disturbances, hot flushes/night sweats, irritability, crawling feelings under the skin) where relevant.

To determine menstrual cycle phase on the day of testing, premenopausal participants collected a urine sample on the morning of their neuromuscular testing session. Enzyme linked immunosorbent assays (ELISA) specific to the urine matrix were used to quantify oestradiol (Abcam, Cambridge, United Kingdom) and progesterone (Invitrogen, Waltham, MA). Urinary hormone levels were used in addition to a menstrual cycle calendar (e.g. day of cycle, last known cycle) to determine menstrual cycle phase using a modified version of Elliott-Sale *et al*. (2021) to account for urinary hormone concentrations.

### Body composition

Dual energy x-ray absorptiometry (DEXA; GE Lunar, Madison, WI) was used to assess body composition, including total body fat and lean mass (kg), and bone mineral density (BMD, g.cm^-2^). During the scan participants lay in a supine anatomical position with the hands in a neutral position. A foam block was placed between the arms and trunk to separate the regions. The enCORE software (GE Healthcare, Chicago, IL) uses skeletal landmarks in the image to detect regions of interest, which were manually adjusted where appropriate. The leg region was determined by placing a border through the femoral neck and the arm region was defined by a border at the medial side of the humerus neck. The sum of leg and arm lean mass was used to calculate appendicular lean mass (kg).

A peripheral quantitative computed tomography (pQCT) scan (voxel size: 0.5 x 0.5mm, scanning speed: 20mm/s, XCT 3000, Stratec Medizintechnik GmBH, Pforzheim, Germany) of the upper thigh was obtained to determine the whole thigh muscle and quadriceps muscle cross-sectional area, and subcutaneous and intramuscular fat area. Participants lay in a supine position with their measured leg secured in a footrest. Tibial length was used as an approximation of femur length, measured from the tibial plateau to the medial malleolus while the knee was flexed to 90° (Cervinka *et al*., 2018). The images were taken at 50% of the tibia length from the mid-condylar cleft towards the hip and analysed using ImageJ software (version 2.0.0; National Institutes of Health, Bethesda, MD, USA). Movement artifacts on the scan were rated from 1 (none) to 5 (extreme) as previously described (Blew *et al*., 2014). Images with a score ≥ 4 were excluded from analysis (*n* = 8). Quadriceps muscle cross-sectional area was calculated by manual tracing using ImageJ software as previously described (Fuchs *et al*., 2023). Whole thigh muscle area and thigh subcutaneous and intramuscular fat were quantified using the pQCT plugin (Rantalainen *et al*., 2011).

### Statistics

All data were analysed using Rstudio 4.3.2 (R Core Team, 2021). To explore the relationship between age and neuromuscular measures, scatterplots were created to visualise the data distribution. For bounded variables (i.e. percentages), beta regression models of the form (*Neuromuscular Measure* ∼ *Age*) were run. Further examination of the scatter plots and residuals of linear models indicated that unbounded variables displayed a non-linear relationship with age. Therefore, generalised additive modelling (GAM) analysis were run with thin plate regression splines and using the Restricted Maximum Likelihood (REML) method to optimise smoothness (Wood, 2017). The model was of the form:

*Neuromuscular Measure* ∼ *s*(*Age*), *method* = *REML*. Model fit was assessed through effective degrees of freedom (EDF) and the k-index (k′), with values closest to 1 indicating linearity and poor fit, respectively. Since the EDF values indicated a non-linear change and visualization of the GAM models suggested a steepness of the slope around the menopausal transition, we identified where this change occurred using the first and second derivatives. The first derivative indicates the slope of the tangent line to the curve, and the second derivative highlights inflexion points. To pinpoint regions of steep decline in the outcome variable, we focused on where the first derivative dropped below 75% of its maximum value.

We then selected the subset of the sample located beyond this threshold to conduct principal component analysis (PCA). The principal components (PCs) were identified and those with eigenvalue above one (i.e., collectively explaining more of the variance than one predictor variable alone) were retained. Correlations between the predictor variables and PCs were visualised in a correlation matrix plot to understand the importance of each variable to the various PCs. We then conducted correlation analysis of the neuromuscular measures with the PCs to identify significant associations. Correlations (r) were classified as strong (|r| ≥ 0.7), moderate (0.7 < |r| ≥ 0.3), weak (|r| < 0.3) or negligible (|r| < 0.1) (Cohen, 1988).

Missing data were imputed using the Multiple Imputation by Chained Equation (mice) package (van Buuren & Groothuis-Oudshoorn, 2011) with predictive mean matching, and the quality of imputation was validated by comparing the distributions of original and imputed data using density and strip plots.

Other packages used in our analysis includes; *FactoMineR* (Lê et al., 2008), *factoextra* (Kassambara, 2020), *mgcv ()* (Wood, 2017)*, lme4* (Bates *et al*., 2015), *lmerTest* (Kuznetsova *et al*., 2017), and *tidyverse* (Wickham *et al*., 2019).

## Results

### Participant characteristics

Eighty-eight biological female participants completed the study, with all decades of adulthood represented in the cohort (18-29 yr: n=19, 30-39 yr: n=13, 40-49 yr: n=12, 50-59 yr: n=16, 60-69 yr: n=15, 70-80 yr: n=13). Participant characteristics including measures of body composition, neuromuscular function, reproductive health and lifestyle components are shown in **Table 1**. Females were recruited at every stage of the reproductive span (premenopausal: n=43, perimenopausal: n=3, postmenopausal: n=42). 41% of pre- or perimenopausal participants used a form of hormonal contraception (IUD, OCP, or implant), and 18% of peri- or postmenopausal participants used hormonal replacement therapy.

**Table 1.**
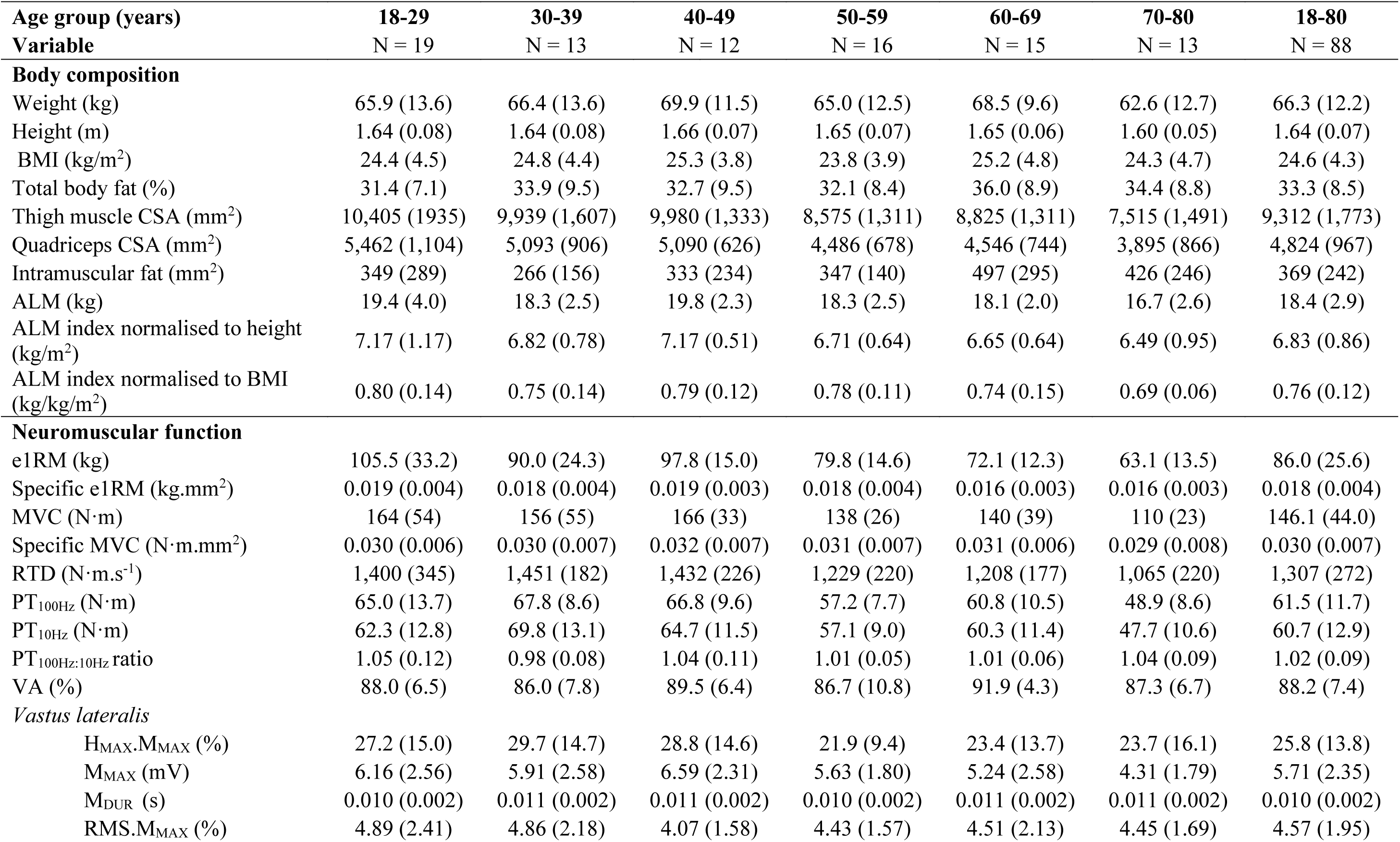
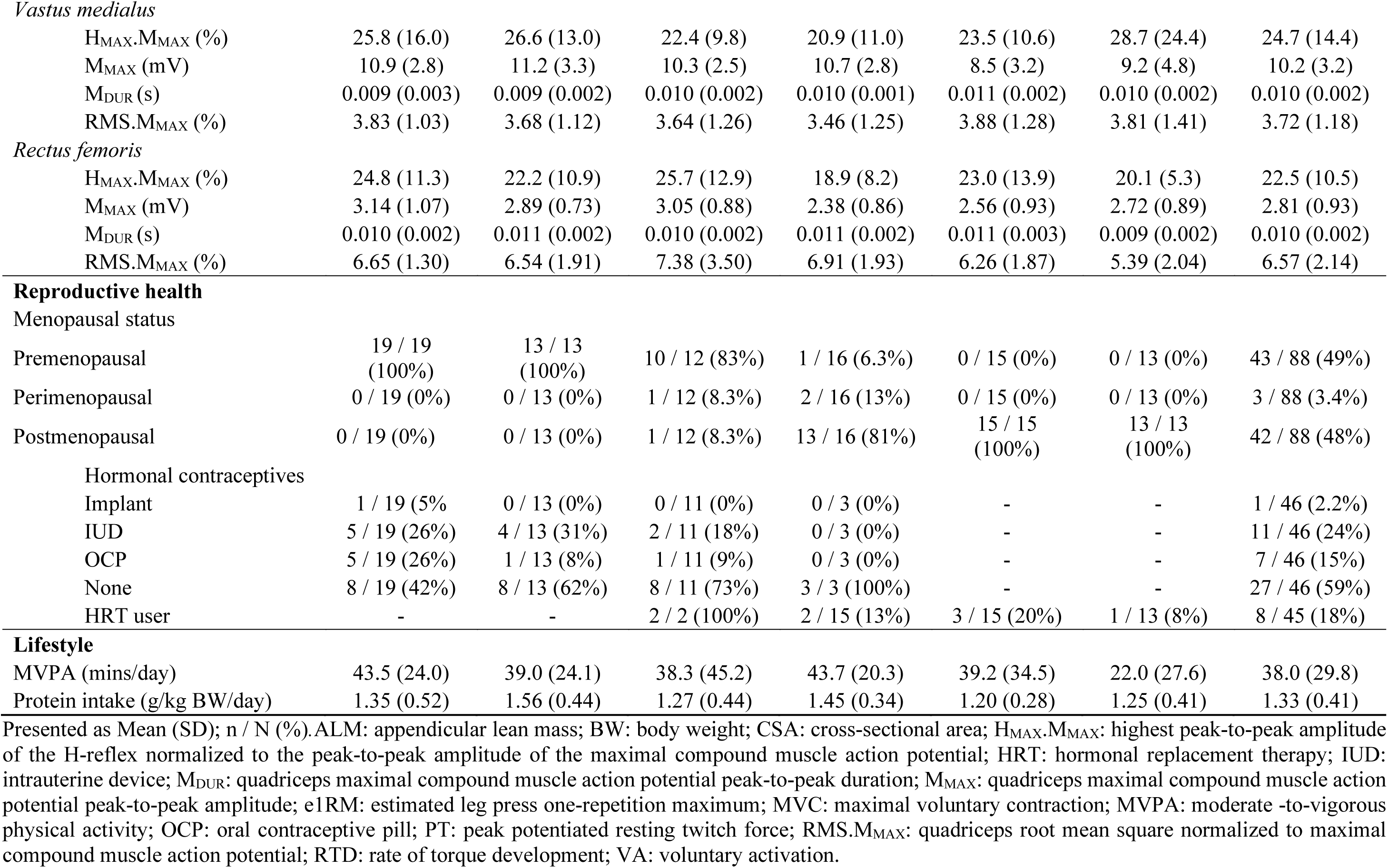
Participant characteristics.

| Age group (years) | 18-29 | 30-39 | 40-49 | 50-59 | 60-69 | 70-80 | 18-80 |
| --- | --- | --- | --- | --- | --- | --- | --- |
| Variable | N = 19 | N = 13 | N = 12 | N = 16 | N = 15 | N = 13 | N = 88 |
| **Body composition** |  |  |  |  |  |  |  |
| Weight (kg) | 65.9 (13.6) | 66.4 (13.6) | 69.9 (11.5) | 65.0 (12.5) | 68.5 (9.6) | 62.6 (12.7) | 66.3 (12.2) |
| Height (m) | 1.64 (0.08) | 1.64 (0.08) | 1.66 (0.07) | 1.65 (0.07) | 1.65 (0.06) | 1.60 (0.05) | 1.64 (0.07) |
| BMI (kg/m<sup>2</sup>) | 24.4 (4.5) | 24.8 (4.4) | 25.3 (3.8) | 23.8 (3.9) | 25.2 (4.8) | 24.3 (4.7) | 24.6 (4.3) |
| Total body fat (%) | 31.4 (7.1) | 33.9 (9.5) | 32.7 (9.5) | 32.1 (8.4) | 36.0 (8.9) | 34.4 (8.8) | 33.3 (8.5) |
| Thigh muscle CSA (mm<sup>2</sup>) | 10,405 (1935) | 9,939 (1,607) | 9,980 (1,333) | 8,575 (1,311) | 8,825 (1,311) | 7,515 (1,491) | 9,312 (1,773) |
| Quadriceps CSA (mm<sup>2</sup>) | 5,462 (1,104) | 5,093 (906) | 5,090 (626) | 4,486 (678) | 4,546 (744) | 3,895 (866) | 4,824 (967) |
| Intramuscular fat (mm<sup>2</sup>) | 349 (289) | 266 (156) | 333 (234) | 347 (140) | 497 (295) | 426 (246) | 369 (242) |
| ALM (kg) | 19.4 (4.0) | 18.3 (2.5) | 19.8 (2.3) | 18.3 (2.5) | 18.1 (2.0) | 16.7 (2.6) | 18.4 (2.9) |
| ALM index normalised to height (kg/m<sup>2</sup>) | 7.17 (1.17) | 6.82 (0.78) | 7.17 (0.51) | 6.71 (0.64) | 6.65 (0.64) | 6.49 (0.95) | 6.83 (0.86) |
| ALM index normalised to BMI (kg/kg/m<sup>2</sup>) | 0.80 (0.14) | 0.75 (0.14) | 0.79 (0.12) | 0.78 (0.11) | 0.74 (0.15) | 0.69 (0.06) | 0.76 (0.12) |
| **Neuromuscular function** |  |  |  |  |  |  |  |
| e1RM (kg) | 105.5 (33.2) | 90.0 (24.3) | 97.8 (15.0) | 79.8 (14.6) | 72.1 (12.3) | 63.1 (13.5) | 86.0 (25.6) |
| Specific e1RM (kg.mm<sup>2</sup>) | 0.019 (0.004) | 0.018 (0.004) | 0.019 (0.003) | 0.018 (0.004) | 0.016 (0.003) | 0.016 (0.003) | 0.018 (0.004) |
| MVC (N·m) | 164 (54) | 156 (55) | 166 (33) | 138 (26) | 140 (39) | 110 (23) | 146.1 (44.0) |
| Specific MVC (N·m.mm<sup>2</sup>) | 0.030 (0.006) | 0.030 (0.007) | 0.032 (0.007) | 0.031 (0.007) | 0.031 (0.006) | 0.029 (0.008) | 0.030 (0.007) |
| RTD (N·m.s<sup>-1</sup>) | 1,400 (345) | 1,451 (182) | 1,432 (226) | 1,229 (220) | 1,208 (177) | 1,065 (220) | 1,307 (272) |
| PT<sub>100Hz</sub> (N·m) | 65.0 (13.7) | 67.8 (8.6) | 66.8 (9.6) | 57.2 (7.7) | 60.8 (10.5) | 48.9 (8.6) | 61.5 (11.7) |
| PT<sub>10Hz</sub> (N·m) | 62.3 (12.8) | 69.8 (13.1) | 64.7 (11.5) | 57.1 (9.0) | 60.3 (11.4) | 47.7 (10.6) | 60.7 (12.9) |
| PT<sub>100Hz:10Hz</sub> ratio | 1.05 (0.12) | 0.98 (0.08) | 1.04 (0.11) | 1.01 (0.05) | 1.01 (0.06) | 1.04 (0.09) | 1.02 (0.09) |
| VA (%) | 88.0 (6.5) | 86.0 (7.8) | 89.5 (6.4) | 86.7 (10.8) | 91.9 (4.3) | 87.3 (6.7) | 88.2 (7.4) |
| *Vastus lateralis* |  |  |  |  |  |  |  |
| H<sub>MAX</sub>.M<sub>MAX</sub> (%) | 27.2 (15.0) | 29.7 (14.7) | 28.8 (14.6) | 21.9 (9.4) | 23.4 (13.7) | 23.7 (16.1) | 25.8 (13.8) |
| M<sub>MAX</sub> (mV) | 6.16 (2.56) | 5.91 (2.58) | 6.59 (2.31) | 5.63 (1.80) | 5.24 (2.58) | 4.31 (1.79) | 5.71 (2.35) |
| M<sub>DUR</sub> (s) | 0.010 (0.002) | 0.011 (0.002) | 0.011 (0.002) | 0.010 (0.002) | 0.011 (0.002) | 0.011 (0.002) | 0.010 (0.002) |
| RMS.M<sub>MAX</sub> (%) | 4.89 (2.41) | 4.86 (2.18) | 4.07 (1.58) | 4.43 (1.57) | 4.51 (2.13) | 4.45 (1.69) | 4.57 (1.95) |
| *Vastus medialis* |  |  |  |  |  |  |  |
| H<sub>MAX</sub>.M<sub>MAX</sub> (%) | 25.8 (16.0) | 26.6 (13.0) | 22.4 (9.8) | 20.9 (11.0) | 23.5 (10.6) | 28.7 (24.4) | 24.7 (14.4) |
| M<sub>MAX</sub> (mV) | 10.9 (2.8) | 11.2 (3.3) | 10.3 (2.5) | 10.7 (2.8) | 8.5 (3.2) | 9.2 (4.8) | 10.2 (3.2) |
| M<sub>DUR</sub> (s) | 0.009 (0.003) | 0.009 (0.002) | 0.010 (0.002) | 0.010 (0.001) | 0.011 (0.002) | 0.010 (0.002) | 0.010 (0.002) |
| RMS.M<sub>MAX</sub> (%) | 3.83 (1.03) | 3.68 (1.12) | 3.64 (1.26) | 3.46 (1.25) | 3.88 (1.28) | 3.81 (1.41) | 3.72 (1.18) |
| *Rectus femoris* |  |  |  |  |  |  |  |
| H<sub>MAX</sub>.M<sub>MAX</sub> (%) | 24.8 (11.3) | 22.2 (10.9) | 25.7 (12.9) | 18.9 (8.2) | 23.0 (13.9) | 20.1 (5.3) | 22.5 (10.5) |
| M<sub>MAX</sub> (mV) | 3.14 (1.07) | 2.89 (0.73) | 3.05 (0.88) | 2.38 (0.86) | 2.56 (0.93) | 2.72 (0.89) | 2.81 (0.93) |
| M<sub>DUR</sub> (s) | 0.010 (0.002) | 0.011 (0.002) | 0.010 (0.002) | 0.011 (0.002) | 0.011 (0.003) | 0.009 (0.002) | 0.010 (0.002) |
| RMS.M<sub>MAX</sub> (%) | 6.65 (1.30) | 6.54 (1.91) | 7.38 (3.50) | 6.91 (1.93) | 6.26 (1.87) | 5.39 (2.04) | 6.57 (2.14) |
| Menopausal status |  |  |  |  |  |  |  |
| Premenopausal | 19 / 19<br>(100%) | 13 / 13<br>(100%) | 10 / 12 (83%) | 1 / 16 (6.3%) | 0 / 15 (0%) | 0 / 13 (0%) | 43 / 88 (49%) |
| Perimenopausal | 0 / 19 (0%) | 0 / 13 (0%) | 1 / 12 (8.3%) | 2 / 16 (13%) | 0 / 15 (0%) | 0 / 13 (0%) | 3 / 88 (3.4%) |
| Postmenopausal | 0 / 19 (0%) | 0 / 13 (0%) | 1 / 12 (8.3%) | 13 / 16 (81%) | 15 / 15<br>(100%) | 13 / 13<br>(100%) | 42 / 88 (48%) |
| Hormonal contraceptives |  |  |  |  |  |  |  |
| Implant | 1 / 19 (5%) | 0 / 13 (0%) | 0 / 11 (0%) | 0 / 3 (0%) | - | - | 1 / 46 (2.2%) |
| IUD | 5 / 19 (26%) | 4 / 13 (31%) | 2 / 11 (18%) | 0 / 3 (0%) | - | - | 11 / 46 (24%) |
| OCP | 5 / 19 (26%) | 1 / 13 (8%) | 1 / 11 (9%) | 0 / 3 (0%) | - | - | 7 / 46 (15%) |
| None | 8 / 19 (42%) | 8 / 13 (62%) | 8 / 11 (73%) | 3 / 3 (100%) | - | - | 27 / 46 (59%) |
| HRT user | - | - | 2 / 2 (100%) | 2 / 15 (13%) | 3 / 15 (20%) | 1 / 13 (8%) | 8 / 45 (18%) |
| **Lifestyle** |  |  |  |  |  |  |  |
| MVPA (mins/day) | 43.5 (24.0) | 39.0 (24.1) | 38.3 (45.2) | 43.7 (20.3) | 39.2 (34.5) | 22.0 (27.6) | 38.0 (29.8) |
| Protein intake (g/kg BW/day) | 1.35 (0.52) | 1.56 (0.44) | 1.27 (0.44) | 1.45 (0.34) | 1.20 (0.28) | 1.25 (0.41) | 1.33 (0.41) |

Presented as Mean (SD); n / N (%).ALM: appendicular lean mass; BW: body weight; CSA: cross-sectional area; H<sub>MAX</sub>.M<sub>MAX</sub>: highest peak-to-peak amplitude of the H-reflex normalized to the peak-to-peak amplitude of the maximal compound muscle action potential; HRT: hormonal replacement therapy; IUD: intrauterine device; M<sub>DUR</sub>: quadriceps maximal compound muscle action potential peak-to-peak duration; M<sub>MAX</sub>: quadriceps maximal compound muscle action potential peak-to-peak amplitude; e1RM: estimated leg press one-repetition maximum; MVC: maximal voluntary contraction; MVPA: moderate -to-vigorous physical activity; OCP: oral contraceptive pill; PT: peak potentiated resting twitch force; RMS.M<sub>MAX</sub>: quadriceps root mean square normalized to maximal compound muscle action potential; RTD: rate of torque development; VA: voluntary activation.

### Effect of the menstrual cycle and hormonal contraception on neuromuscular outcomes in premenopausal females

This study is part of a larger study relying on the collection of muscle biopsies during the early follicular phase of the menstrual cycle in premenopausal females (and in perimenopausal females, where possible), or at any time in postmenopausal females. As a result, and while all efforts were made to collect neuromuscular data during the same menstrual cycle phase, this was not always possible. As a first step, we verified whether menstrual cycle phases had a significant effect on any of the neuromuscular outcomes. All measures but two (rectus femoris M_MAX_ and vastus medialis M_DUR_, both p < 0.05) displayed no significant association with menstrual cycle phase (**Table 2**). The same analysis was run to investigate the effect of hormonal contraception on neuromuscular outcomes in pre- and perimenopausal females. There was no significant association between hormonal contraception mode (none, implant, IUD or OCP) and any of the neuromuscular outcomes (**Table 2**).

**Table 2.**
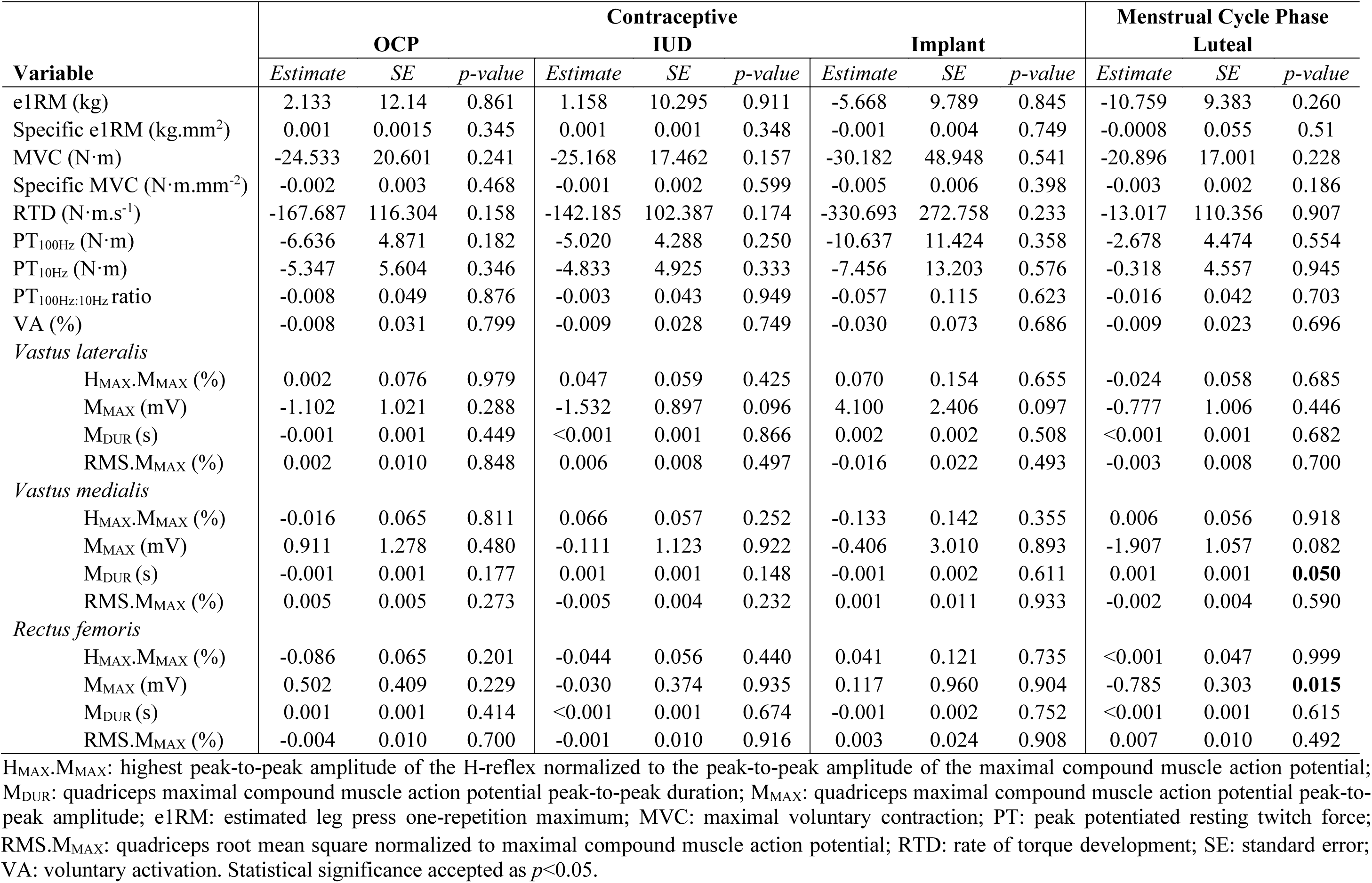
Effects of hormonal contraception and menstrual cycle phase on neuromuscular outcomes.

### Effect of age on neuromuscular outcomes

Beta-regression models were conducted for bounded neuromuscular outcomes (i.e., outcomes expressed as percentages) including voluntary activation and quadriceps H-reflex and RMS.M_MAX_. None of these variables were significantly associated with age (**Table 3**).

**Table 3.**
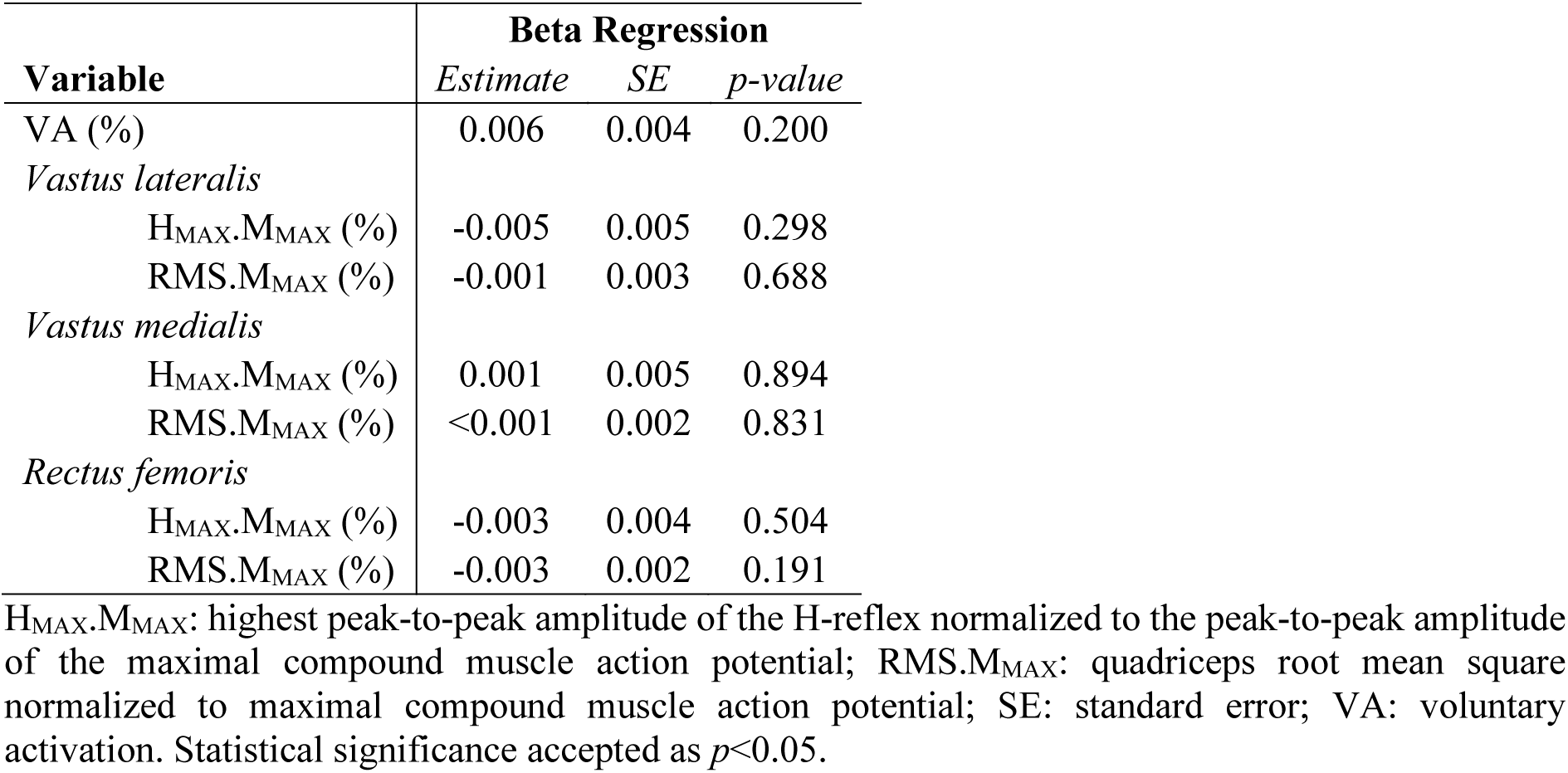
Effect of age on bounded neuromuscular outcomes using beta-regression.

All other outcomes were modelled using generalized additive models (GAM). The smooth terms for MVC (N·m), e1RM (kg), RTD (N·m.s^-1^), PT_100_ (N·m) and PT_10_ (N·m) indicated a declining non-linear relationship with age (p < 0.001; **Figure 1A-E**), meaning that, as age increased, these outcomes decreased in a non-linear manner. RFM_MAX_ (mV), quadriceps lean CSA (mm²) and e1RM (kg.mm^2^) were associated with age in a negative, quasi-linear manner (p < 0.05, **Figure 1F, G, I**). No other outcomes were significantly associated with age (**Table 4**) including MVC (N·m.mm²) (**Figure 1H**).

**Figure 1:**
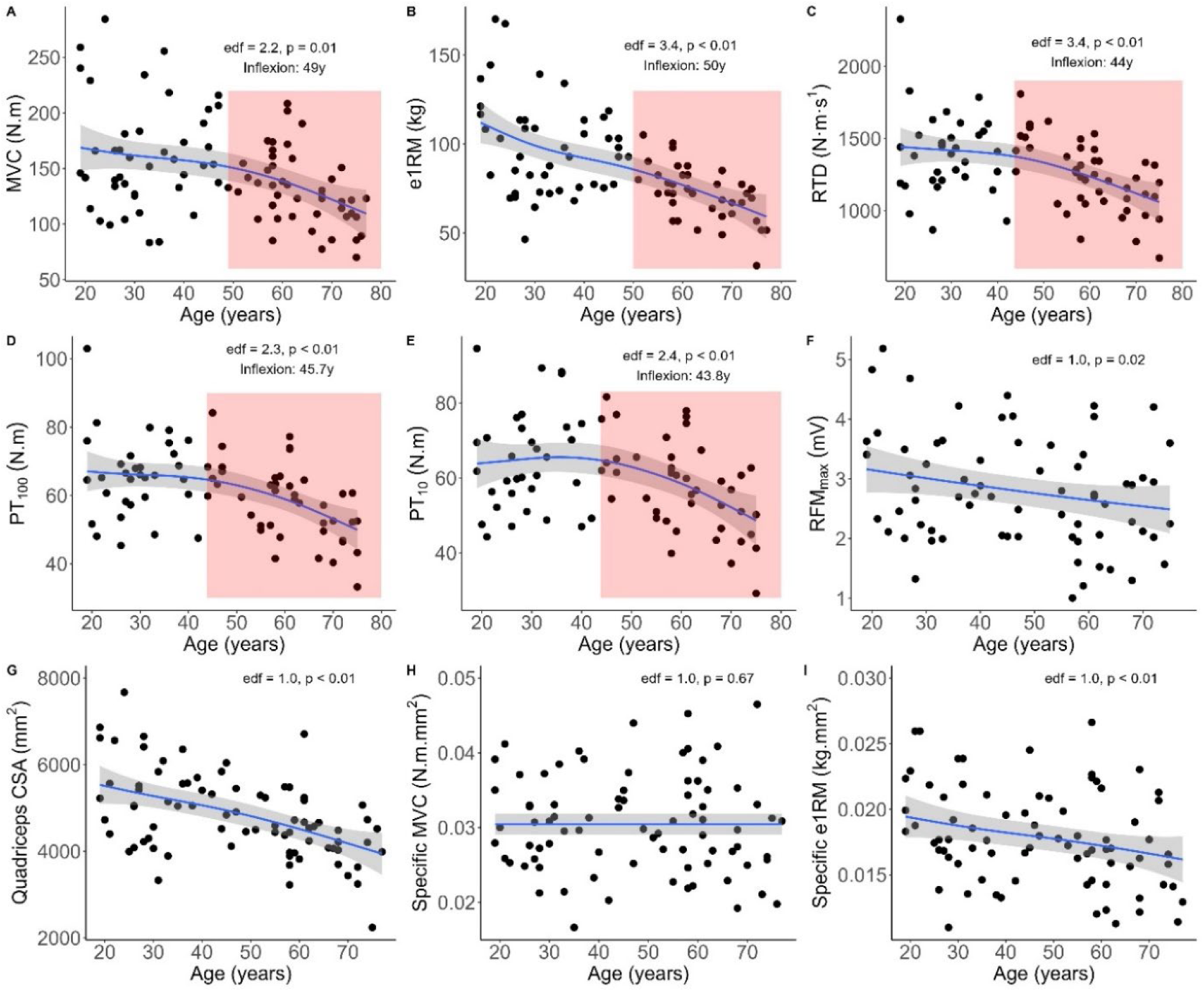
Associations between neuromuscular outcomes and age. All neuromuscular outcomes were modelled using generalized additive models (GAM). The red shaded region highlights a rapid decline in the outcome, starting at the point its first derivative falls below the 75% threshold. edf = estimated degree of freedom. p < 0.05 were considered statistically significant.

**Table 4.** Effect of age on neuromuscular outcomes using generalised additive models.

| Variable | GAM Models |  |
| --- | --- | --- |
|  | EDF | p-value |
| e1RM (kg) | 3.389 | **<0.001** |
| Specific e1RM (kg.mm<sup>2</sup>) | 1.000 | **0.007** |
| MVC (N·m) | 2.196 | **0.001** |
| Specific MVC (N·m.mm<sup>2</sup>) | 1.001 | 0.673 |
| RTD (N·m.s<sup>-1</sup>) | 2.005 | **<0.001** |
| PT<sub>100Hz</sub> (N·m) | 2.272 | **<0.001** |
| PT<sub>10Hz</sub> (N·m) | 2.410 | **0.001** |
| PT<sub>100Hz:10Hz</sub> ratio | 2.164 | 0.386 |
| *Vastus lateralis* |  |  |
| M<sub>MAX</sub> (mV) | 1.294 | 0.107 |
| M<sub>DUR</sub> (s) | 1.000 | 0.212 |
| *Vastus medialis* |  |  |
| M<sub>MAX</sub> (mV) | 1.000 | 0.076 |
| M<sub>DUR</sub> (s) | 1.000 | 0.400 |
| *Rectus femoris* |  |  |
| M<sub>MAX</sub> (mV) | 1.000 | **0.019** |
| M<sub>DUR</sub> (s) | 1.711 | 0.494 |
| Quadriceps CSA (mm<sup>2</sup>) | 1.001 | **<0.001** |

EDF: effective degrees of freedom; M<sub>DUR</sub>: quadriceps maximal compound muscle action potential peak- to-peak duration; M<sub>MAX</sub>: quadriceps maximal compound muscle action potential peak-to-peak amplitude; e1RM: estimated leg press one-repetition maximum; MVC: maximal voluntary contraction; PT: peak potentiated resting twitch force; RTD: rate of torque development. Statistical significance accepted as $p < 0.05$ .

The fitted GAM models that were significant for age and displayed a significant EDF value greater than 1 were selected for further analysis. The steeper decline of each curve, defined as the point where the first derivative fell below 75% of its maximum value, started at 43.7 years of age for RTD (N·m.s^-1^), 43.8 years of age for PT_10_ (N·m), 45.7 years of age for PT_100_ (N·m), 49 years for MVC (N·m) and 50 years of age for e1RM (kg) (**Figure 1A-E**, red shaded region).

### Post menopause analysis

PCA analysis was conducted on the subset of the sample aged beyond the relevant inflexion points to investigate the potential contribution of body composition, physical activity and sex hormones to the observed decline of neuromuscular outcomes postmenopause. Principal components 1 to 4 accounted for 70% of the total variance in the data (PC1 = 24%; PC2 = 20%; PC3 = 14%; PC4 = 12%). Each of these components had eigenvalues greater than 1, indicating that they explained more variance than any individual variable, and thus, were considered significant in capturing the underlying patterns of the dataset. The significant variables within each component differed between each PC (**Figure 2a**). Correlation of the PCs with neuromuscular variables that were significantly affected by age (e1RM (kg), MVC (N·m), RTD (N·m.s-1), PT10 (N·m) and PT100 (N·m)) were then examined. PC1 was correlated with all variables, PC3 was correlated with PT_10_ (N·m) and PT_100_ (N·m), and PC4 was correlated with all variables other than e1RM (**Figure 2b**).

**Figure 2:**
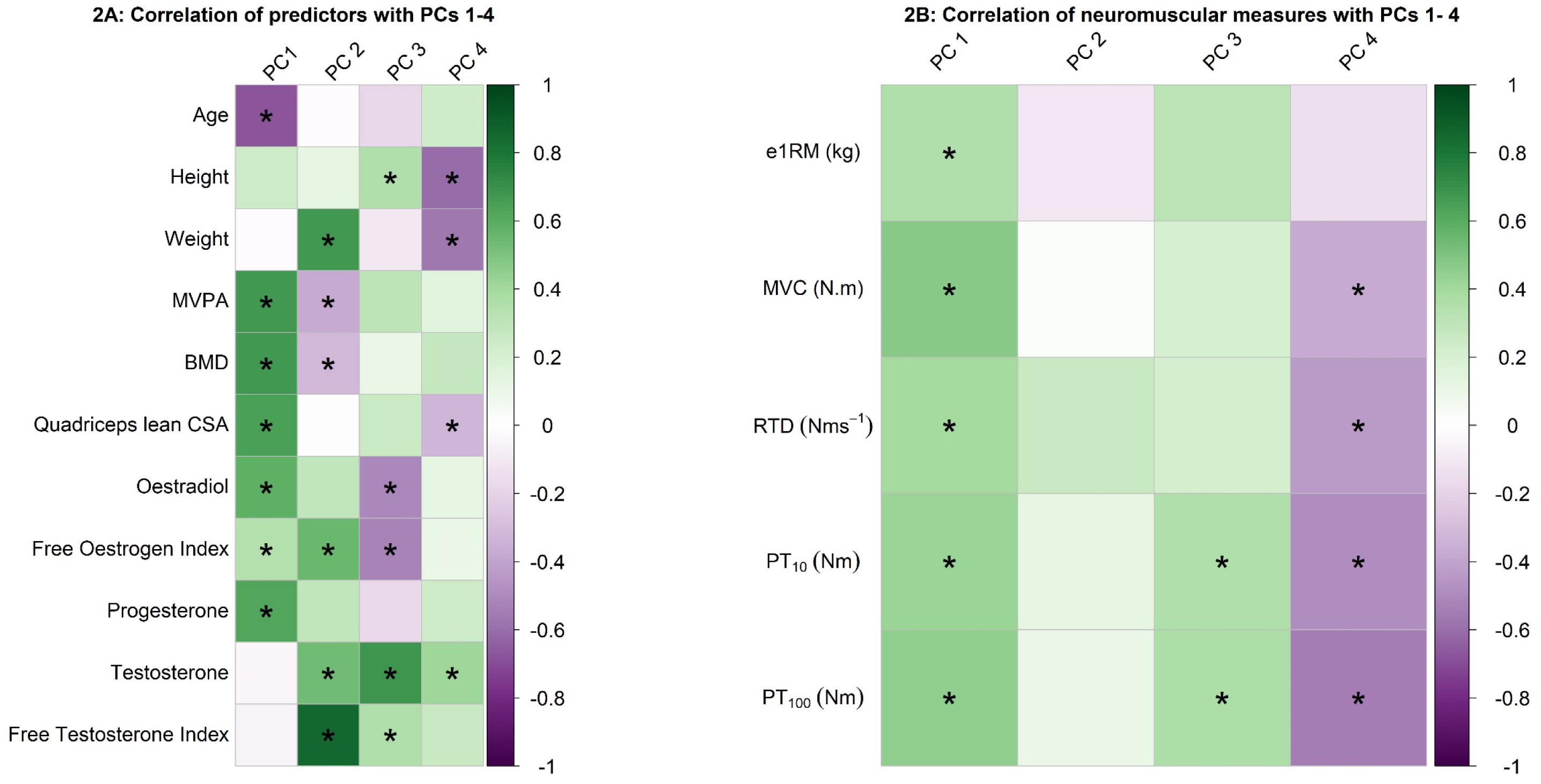
PCA analysis of Postmenopausal Females. **A)** Correlogram: Correlations between the predictor variables and the principal components. **B)** Correlogram: Correlations between neuromuscular outcomes and principal components. Negative correlations are shown in purple, and positive correlations are shown in green. *Indicates a significant correlation at p < 0.05.

## Discussion

This study aimed to investigate the determinants of age-related changes in quadriceps neuromuscular function across the adult female lifespan. Analysis of 88 healthy females represented equally across each decade of life from 18-80 years demonstrated a non-linear decline and accelerated reductions occurring around menopausal onset in all isometric voluntary and evoked torque responses, whereas quadriceps lean CSA and rectus femoris compound muscle action potential amplitude decreased quasi-linearly with age. No effect of age was observed for isometric MVC normalized to quadriceps lean CSA, although e1RM relative to quadriceps lean CSA decreased. Outcomes representative of excitatory neural drive to quadriceps were unaffected by age, including voluntary activation, H-reflexes, and RMS of vastus lateralis, vastus medialis and rectus femoris normalized to maximal muscle compound action potential. PCA analysis revealed that, in postmenopausal females, age-related reductions in neuromuscular variables could be explained by inter-individual differences in body composition, moderate to vigorous physical activity level and concentrations of sex hormones oestrogen, progesterone and testosterone.

### Effect of Age on Neuromuscular Function

#### Voluntary and Evoked Torques

Non-linear age-related reductions in muscle torque starting in the fourth decade of the female lifespan aligns with previous observations of muscle strength (Lindle *et al*., 1997) and general molecular dysfunction within several physiological systems (Shen *et al*., 2024). Muscle torque is highest when shortening velocity is lowest (Thorstensson *et al*., 1976) and torque generating capacity is directly proportional to the CSA of a muscle (Jones *et al*., 2008). The lack of age-related change in isometric voluntary torque relative to quadriceps lean CSA supports the considerable role that muscle size/hypertrophy plays in isometric torque production across the female lifespan. In the present study, quadriceps lean CSA decreased quasi-linearly at an average rate of ∼29mm² per year, slightly less than the linear decrease reported from 40 years of age for healthy females (Mizuno *et al*., 2021). However, the decline in isometric MVC was non-linear with an inflexion point reported at 49 years of age, suggesting that quadriceps lean CSA becomes relatively more important for isometric torque production beyond this age. Maintenance of skeletal muscle mass later in life has vital implications for metabolic and musculoskeletal health and the prevention of age-related diseases such as type 2 diabetes, sarcopenia and osteoporosis (Evans, 1997; Lombardi *et al*., 2016). Although the lack of change in isometric MVC relative to quadriceps lean CSA aligns with some (Laakkonen *et al*., 2017), others have shown a decrease in isometric muscle specific torque with increasing age (Wrucke *et al*., 2024). The reason for this discrepancy is unclear but may be related to methods adopted to quantify quadriceps CSA. Here, we used computed tomography and extracted lean cross-sectional area from the quadriceps independent of non-contractile tissue (intramuscular and subcutaneous), which is a validated technique with high level of agreement with gold standard magnetic resonance imaging methods (Fuchs *et al*., 2023).

Discordant with the physiological determinants of isometric torque with advancing age, quadriceps type II fibres preferentially atrophy in older age (Nilwik *et al*., 2013) and explain a large proportion of the age-related decline in dynamic strength and power (Power *et al*., 2013). Moreover, loss of high threshold motor units in older adults decreases motor unit firing frequencies (Piasecki *et al*., 2016; Wages *et al*.) and older females show less musculotendinous stiffness than their young counterparts (Wu *et al*., 2016). Indeed, muscle size/hypertrophy plays less of a role in e1RM compared to isometric MVC, as significant decreases across the lifespan remained evident when normalized to quadriceps lean CSA. Although dynamic torque and muscle shortening velocity decreases earlier in the lifespan and at greater magnitude compared to torque production at slower speeds (Raj *et al*., 2010; Haynes *et al*., 2020), we did not observe any differences in the age of decline for isometric MVC compared to dynamic e1RM. Quadriceps activation is similar during isometric knee extension and leg press exercise (Alkner *et al*., 2000) and knee extension joint power can effectively model torque-velocity relationships in leg press (Bobbert, 2012), demonstrating the significant contribution of the quadriceps to leg press performance. However, the lack of difference in the age of decline for isometric MVC and dynamic leg press may have been confounded by contribution of other large muscle groups which also demonstrate near maximal activation levels during leg press exercise at heavy loads (e.g., gluteus maximus) (Da Silva *et al*., 2008) or from potential influence of the muscle stretch-shortening cycle from a repetitive 5RM test (Cormie *et al*., 2011). Nonetheless, reductions in dynamic torque and power have greater negative consequences for overall functional capacity (Power *et al*., 2013) and are more prevalent in older females compared to males (Wrucke *et al*., 2024).

Interestingly, the inflexion point of accelerated age-related reductions in MVC and e1RM (49-50 years) occurred after the observed reduction in PT_10_, RTD (both 44 years) and PT_100_ (46 years). Although PT_10_ seemed to reduce slightly earlier in the lifespan than PT_100_, the lack of change in the high to low frequency ratio suggests a comparable rate of impairments in different peripheral compartments. Considerable reductions in high and low frequency evoked torque outcomes have previously been shown in comparisons between younger and older females (Solianik *et al*., 2017; Varesco *et al*., 2022; Wrucke *et al*., 2024), however, here we illustrate the specific age of which marked reductions start to occur around menopause onset during the fourth decade. The reduced peak torque during high frequency stimulation may indicate neuromuscular junction instability arising from age-related morphological alterations (e.g., number and affinity of acetylcholine receptors, reduced junctional folds and increased fragmentation) (Hepple & Rice, 2016; Piasecki *et al*., 2016; Arnold & Clark, 2023) or increased sarcolemma refractoriness related to sodium channel inactivation (Lee *et al*., 2018). Reduced peak torque during low frequency stimulation suggests age-related impairments in Ca^2+^ handling/content (Jones, 1996; Millet *et al*., 2011) known to occur in elderly females (Hunter *et al*., 1999) and related to reduced maximal rate of torque development and strength of actin-myosin bound states in males (Mazara *et al*., 2021). The earlier onset of age-related impairments in evoked compared to voluntary torque responses suggests that intrinsic muscle function may be impaired earlier in the lifespan than any observable decline in functional capacity, perhaps related to compensatory increases in motor unit firing frequencies resulting from reduced muscle quality. Although poor muscle quality is suggested to in part explain higher motor unit firing rates in older females compared to males (Guo *et al*., 2024), this requires further investigation within females and during high force contractions as our results show whole thigh intramuscular fat did not increase until the sixth decade, whereas others show increased intermuscular fat infiltration in quadriceps from the fourth decade (Mizuno *et al*., 2021).

#### Voluntary Activation and Electromyography

The lack of age effect on quadriceps voluntary activation, H-reflex amplitude and RMS normalized to maximal muscle compound action potential suggests that steep reductions in quadriceps torque and neuromuscular function starting during the fourth decade and during the menopausal transition were largely mediated by mechanisms occurring at or distal to the neuromuscular junction rather than an incapacity of supraspinal and spinal circuits to volitionally recruit existing quadriceps motor units or reduced facilitation of the motoneuron pool via Ia afferent stimulation. Previous studies show that compared to younger cohorts, quadriceps voluntary activation is lower in older females when assessed by train stimuli (Solianik *et al*., 2017), decreased in older mixed-sex cohorts with high-frequency paired pulse (Mau-Moeller *et al*., 2013), or unchanged when assessed via transcranial magnetic stimulation (Wrucke *et al*., 2024) or magnetic stimulation (Varesco *et al*., 2022). Here, we implemented gold-standard technique to assess voluntary activation of large functional quadriceps muscles in females via high frequency doublet twitch interpolation (Nuzzo *et al*., 2019) and normalized voluntary EMG RMS amplitude to maximal M-wave to account for muscle size and peripheral transmission confounding effects (Millet *et al*., 2011). The heterogeneity in current findings on voluntary activation in ageing females may be related to assessment method (Nuzzo *et al*., 2019; Rozand *et al*., 2020), although a lack of decrease in voluntary activation and RMS with age in the present study may be related to physical activity level of the included participants. All participants in the present study including those above 65 years met (or exceeded for those 50-59 years) the minimum guidelines for moderate to vigorous physical activity (150 – 300min per week) (Bull *et al*., 2020), reflecting a well-known recruitment bias in exercise physiology studies (Stefanetti *et al*., 2014). Long-term physical activity may promote reinnervation of denervated fibres (Piasecki *et al*., 2019) without impacting muscle size or isometric and dynamic torque when the physical activity is aerobic in nature (Chambers *et al*., 2020).

Rectus femoris M-wave amplitude decreased with increasing age without any observed changes for vastii muscles, suggesting potential vulnerability of this bi-articular quadriceps muscle to age-related impairments in neuromuscular function across the female lifespan. Rectus femoris typically demonstrates higher percentage of MyHC type II fiber isoforms compared to mono-articular vastii (∼62 vs. 54%) (Johnson *et al*., 1973) and these fibers preferentially atrophy into older age (Andersen, 2003; Hepple & Rice, 2016; Hunter *et al*., 2016). Further, sodium channel inactivation in rectus femoris in older females has been attributed to downregulation in Na^+^/K^+^ ATPase (Lee *et al*., 2018) contributing to potential denervation (Clausen, 2003), although at a single muscle fibre level in vastus lateralis, α1 and α2 Na^+^/K^+^ ATPase isoforms are not different between older and younger mixed-sex cohorts (Wyckelsma *et al*., 2016). Although we could not reliably segment different quadriceps lean muscle cross-sectional areas, Mizuno *et al*. (2021) manually segmented quadriceps muscles (inclusive of lean and intermuscular fat mass) from computed tomography and reported comparable inter-muscular reductions in whole cross-sectional area beyond 40 years of age in healthy females. Thus, reduced M-wave amplitude for rectus femoris is likely explained by a contribution of both reductions in lean muscle cross-sectional area and denervation. Indeed, denervation occurs prior to muscle atrophy due to axonal sprouting and motor unit remodelling (Hunter *et al*., 2016). Rectus femoris activation during locomotor tasks (e.g. cycling, walking or jogging) of moderate to high intensity are much lower compared to vastii, with recruitment levels > 90% MVC reached only during maximal intensity (O’Bryan *et al*., 2018), suggesting maximal intensity exercise may be necessary to mitigate rectus femoris neuromuscular degeneration in older age. Rectus femoris plays a significant role in torque generation and transfer across the hip and knee joints during locomotor functional tasks (Pandy & Andriacchi, 2010), illustrating its importance in functional capacity of older adults. Although more investigation is warranted to dissociate rectus femoris denervation/atrophy from an increase in muscle-electrode distance arising from age-related increases in subcutaneous and intramuscular fat mass (Nordander *et al*., 2003; Akima *et al*., 2015; Mizuno *et al*., 2021), we observed no decrease in vastii M-wave amplitudes despite others reporting greater age-related increases in vastus lateralis adipose tissue area compared to rectus femoris (Akima *et al*., 2015). Potential confounding effects of menstrual cycle phase on rectus femoris M_MAX_ also warrants further investigation to validate our interpretation.

### Physical Activity, Body Composition and Hormonal Influence Post menopause

Principal component #1 explained 24% of the variance in the data and was correlated with all significant neuromuscular outcomes. This dimension included strong positive correlations with MVPA and bone density; moderate positive correlations with quadriceps lean CSA, progesterone, oestrogen and free oestrogen index (i.e. bioactive and not bound to SHBG); and a strong negative correlation with age. Aside from the well described negative consequences of advancing age on isometric and dynamic strength, this finding illustrates the interplay between physical activity, muscle mass, bone mass and primary female sex hormones on the capacity for maximal isometric and dynamic torque production. Skeletal muscle and bone are tightly interconnected, with internal forces stimulating dose-dependent changes in bone formation (Rubin & Lanyon, 1984) and various skeletal muscle myokines and growth factors (e.g., insulin-like growth factor-1, interleukin-6), which influence muscle protein synthesis and bone turnover rate (Gomarasca *et al*., 2020). Further, musculoskeletal structure and function are both sensitive to oestrogens (Almeida *et al*., 2017; Alexander *et al*., 2022) and physical activity (O’Bryan *et al*., 2022). At physiological concentrations, we reported that progesterone may promote muscle protein synthesis and oestrogen may improve muscle contractile function (Alexander *et al*., 2022; Critchlow *et al*., 2023). Others have shown oestrogen and progesterone replacement therapies may be beneficial to voluntary muscle strength (Greising *et al*., 2009) but not muscle mass (Javed *et al*., 2019), suggesting potential positive effects of these sex hormones on skeletal muscle quality and/or motor unit recruitment.

Approximately 14% of the variance in data was explained by principal component #3, which was significantly correlated with PT_100_ and PT_10_. Aside from a positive correlation with height (known to correlate with femur length and increase torque production), principal component #3 was negatively correlated with total and free oestrogen and positively correlated with total and free testosterone, suggesting that individuals with lower E2 but higher androgen levels after menopause have a mitigated loss of muscle function arising from physiological mechanisms occurring at or distal to the neuromuscular junction. Thus, this finding may suggest a potential compensatory mechanism whereby low oestradiol may be counteracted by increasing androgens to maintain intrinsic muscle function. Indeed, declining ovarian function during menopause decreases the circulating concentrations of multiple hormones, but especially oestradiol, increasing the testosterone to oestradiol ratio in both total and bioavailable (i.e., not bound to SHBG) forms (Burger *et al*., 2008; Pöllänen *et al*., 2011). The relative importance of free oestrogen and androgens is further highlighted by principal component #2, which explained 20% of the variance, although this was not correlated to any of the measured neuromuscular variables. Thus, bioavailable forms of these hormones may influence underlying neuromuscular mechanisms not captured by the present study, including supraspinal or spinal circuits (Piasecki *et al*., 2024), or molecular processes within skeletal muscle fibres (Alexander *et al*., 2024). Overall, further investigation into the role of different forms of primary sex hormones in neuromuscular ageing across the female lifespan is warranted.

Finally, principal component #4 explained 12% of the variance and was associated with reductions in isometric voluntary and evoked torques but not dynamic e1RM, and was explained by lower measures of body composition (height, weight and quadriceps lean CSA) and higher levels of total testosterone. These findings support the role of quadriceps lean CSA and overall muscle mass on capacity for isometric voluntary and evoked torque production. Although reduced isometric torque was associated to higher total testosterone, total testosterone is consistently shown to have minimal effect on lean mass or muscle strength in pre (Alexander *et al*., 2021; Alexander *et al*., 2024) and postmenopausal females (Alexander *et al*., 2022).

### Menstrual Phase and Contraception Effects

Previous evidence suggests that some measures of neuromuscular function may fluctuate with hormonal changes across the menstrual cycle. Downstream at the spinal motoneurons, presynaptic inhibition of Ia sensory afferents (Hoffman *et al*., 2018) and firing rate of vastus lateralis motor units (Piasecki *et al*., 2023) are reduced during the ovulation to mid-luteal phase of the menstrual cycle. Premenopausal females also display higher persistent inward currents in several lower-limb muscles compared to young males, suggesting a potential role of sex hormones in neuromodulation of spinal motoneurons (Jenz *et al*., 2023). While some have reported potential menstrual cycle effects on corticospinal and intracortical circuits (Ansdell *et al*., 2019), others have shown limited effect on skeletal muscle contractile function (de Jonge *et al*., 2001) and most studies report negligible effects of the menstrual cycle or contraceptive use on isometric or dynamic strength performance (McNulty *et al*., 2020). Thus, although the phase and contraceptive effects reported here are between participants, we suspect no or trivial effects occurred within participants.

## Conclusions

This study is the first to evaluate female neuromuscular function evenly across each decade from 18 to 80 years of age and to attempt to associate any observed decline with inter-individual differences in sex hormone concentrations. The onset of neuromuscular degeneration started during the fourth decade and menopause onset, with intrinsic muscle function reducing before any observed functional decline. Reductions in isometric voluntary torque were mostly explained by changes in muscle mass, whereas reductions in dynamic torque during leg press were more complex. Neuromuscular degeneration in ageing females may be more pertinent for rectus femoris than for other quadriceps muscles, suggesting potential vulnerability to ageing and functional impairment. Although some of the variance in the postmenopausal data was explained by differences in body composition and physical activity level, all principal components involved significant correlations with total or free sex hormone concentrations (oestrogen, progesterone and/or testosterone), illustrating the potential importance of these primary sex hormones in neuromuscular function in ageing females. Thus, future research should explore how interventions aimed at addressing dramatic and sudden decreases in sex hormone concentrations around menopause onset may help mitigate age-related neuromuscular degeneration across the female lifespan.

## Data availability statement

The R code for analysis can be found https://github.com/DaniHiam/NeuroMusc_FAME. Raw data is available upon reasonable request.

## Author Contributions

SOB was involved in conception and experimental design, acquisition, analysis and interpretation of data, and drafting and revising the manuscript for intellectual content. AC was involved in acquisition, analysis and interpretation of data, and drafting and revising the manuscript for intellectual content. CF was involved in analysis of data and revising the manuscript for intellectual content. DH was involved in conception and experimental design, acquisition, analysis and interpretation of data and drafting and revising the manuscript for intellectual content. SL was involved in conception and experimental design, interpretation of data and drafting and revising the manuscript for intellectual content. All authors approved the final version to be published and agree to be accountable for all aspects of the work.

